# A Phospho-Switch for Cell Fate Control

**DOI:** 10.64898/2026.02.21.707151

**Authors:** Jin Ming, Xianzhuang Liu, Zexiao Jia, Wei Shi, Jiajun Li, Shikun Wang, Yulin Chen, Shixian Lin, Yu Liang, Peng Guo, Hanqing Zhao, Yuxiang Yao, Ruona Shi, Xiaofei Zhang, Yuanyue Shan, Yu Fu, Bo Wang, Chengchen Zhao, Duanqing Pei

## Abstract

Cell fate control is thought to be complex, likely involving both cell intrinsic and extrinsic factors, and remains ill-defined at the molecular level. Here we show a phospho-switch controls cell fate both in vitro and in vivo. We first show that SALL4 is phosphorylated at multiple sites, but only T903 as the primary one as SALL4^T903A^ lost BAF interaction and ∼90% activity in reprogramming. Mechanistically, we demonstrate that SALL4^pT903^ is sensitive to BMP4 signaling through DUSP9 axis. Finally, we show by tetraploid embryo complementation that mESCs harboring SALL4^T903A^ can sustain embryo development but with severe defects including cranial hypoplasia and flattened skull vault after birth. A genome wide search identifies 608 transcription factors harboring the same HTG motif sandwiched by two zinc fingers, suggesting that this phospho-switch may be a conserved mechanism to control cell fate.

**In brief:** A phosphorylation switch, pT903, in SALL4 governs cell fate by modulating BAF complex interaction in response to BMP4-DUSP9 signaling, as its dephosphorylation abrogates reprogramming and causes cranial defects in mice, suggesting that such switches may broadly regulate cell fate machinery.

**Highlights:**

- A T903 phospho-switch in SALL4 gates BAF interaction to orchestrate gene activation and reprogramming
- BMP4-DUSP9 signaling converges on SALL4^pT903^ dephosphorylation to couple extracellular cues with cell fate determination
- Disrupting the SALL4-T903 phospho-switch causes severe postnatal developmental defects in mice.
- The HTGE motif may function as a central switch on key transcription factors for sensing upstream signals

## INTRODUCTION

Multicellular organisms are constructed in such a way that each cell is generated with precision to execute its designated function^1^. How such precision is encoded in the genome and decoded by a cell is one of the most fascinating questions in modern biology. Insights generated from model organisms such as *Drosophila* have illuminated the link between specific genetic loci in the genome to morphogenic events in development^2^. The best example is the homeobox genes that specify the body plan of *Drosophila*^3,4^, delineating a logic universal across species examined so far, suggesting that information hardwired in the genome dictates the precision in development. At the cellular and molecular level, it has been well characterized that homeobox genes are transcription factors (TFs) that control the expression of target genes which then orchestrate the precise generation and spatial locations of each cell to endow a tissue or organ in precision^5,6^.

Thus, how TFs control the fate of newly generated cells has been a focus in recent years^7^.

The problem of cell fate control or CFC has been approached experimentally both in vitro and in vivo. In vivo lineage tracing has generated enormous datasets to illuminate the diversity of cells from a common progenitor, resulting in an understanding of cell fate decision with unprecedented resolution at single cell level^8^. In vitro studies should complement in vivo lineage tracing to yield more precision model of cell fate control at the molecular level as more sophisticated methods can be deployed to determine the precise molecular mechanism associated with cell fate control. Indeed, the reprogramming paradigm has provided key insights into cell fate control^9^ such as the role of epigenetic complexes such as Sin3A (Switch-Independent 3A)^10,11^, NuRD (Nucleosome Remodeling and Deacetylase complex)^12–14^ and BAF (Brg/Brm-associated factors complex)^15–17^ in guiding somatic cells such as E13.5 (embryo day 13.5) MEFs (mouse embryonic fibroblast cells) to pluripotent mESCs (mouse embryonic stem cells) equivalent or iPSCs (induced pluripotent stem cells). These efforts have led to a paradox that the same factors that orchestrate normal developmental processes (forward) often are reprogramming (backward) factors to reverse cell fate from MEFs to iPSCs.

To reconcile this paradox, we hypothesized that each cell contains a cell intrinsic machine or CiM or cell fate machine or CFM. This machine drives cell fate decision regardless of being forward or backward as long as a cell interprets the hardwired information in the genome based on cues both intrinsic and extrinsic. In this report, we provide a critical component of this mechanism by identifying a phospho-switch encoded at threonine/T903 of SALL4, a homolog of *Drosophila* region specific homeotic gene *spalt,* that regulates mouse cell fate control both in in-vivo development and in-vitro reprogramming.

## RESULTS

### Phosphorylation at T903 is critical for SALL4 function

To elucidate how a single transcription factor orchestrates diverse cell fate decisions both in vivo and in vitro, our attention has been directed toward a unique transcription factor, SALL4. Functioning critically in development, disease, aging, and cellular reprogramming across species, SALL4 has been documented to interact with a range of macromolecular machineries with distinct functionalities. Nonetheless, the regulatory mechanisms governing its functional switching remain unreported to date.

To investigate the molecular mechanisms underlying SALL4 function, we leveraged its established role in the JGES (*Jdp2*-*Glis1*-*Esrrb*-*Sall4*) reprogramming system, which converts E13.5 mouse embryonic fibroblasts (MEFs) into induced pluripotent stem cells (iPSCs)^14^, and performed immunoprecipitation-mass spectrometry (IP-MS) (Figure 1A). We searched the secondary spectral library for phosphorylation, acetylation, and SUMOylation modifications on SALL4. The results show that only phosphorylation can be detected. Specifically, we have identified eight phosphorylated amino acid residues, S122, S135, T410, S509, S760, S785, S798, and T903. Among these, all except S785 and S798 are novel phosphorylation sites^18,19^(Figure 1B, 1C and S1A). To assess the functional relevance of these modifications, we generated alanine-substitution mutants at each phosphorylated residue to abolish phosphorylation. All mutants exhibited reduced reprogramming efficiency compared to wild-type SALL4. The T903A mutant showed the most severe defect, lost ∼90% activity in reprogramming (Figure 1D), underscoring the critical role of pT903 in SALL4.

**Figure 1.**
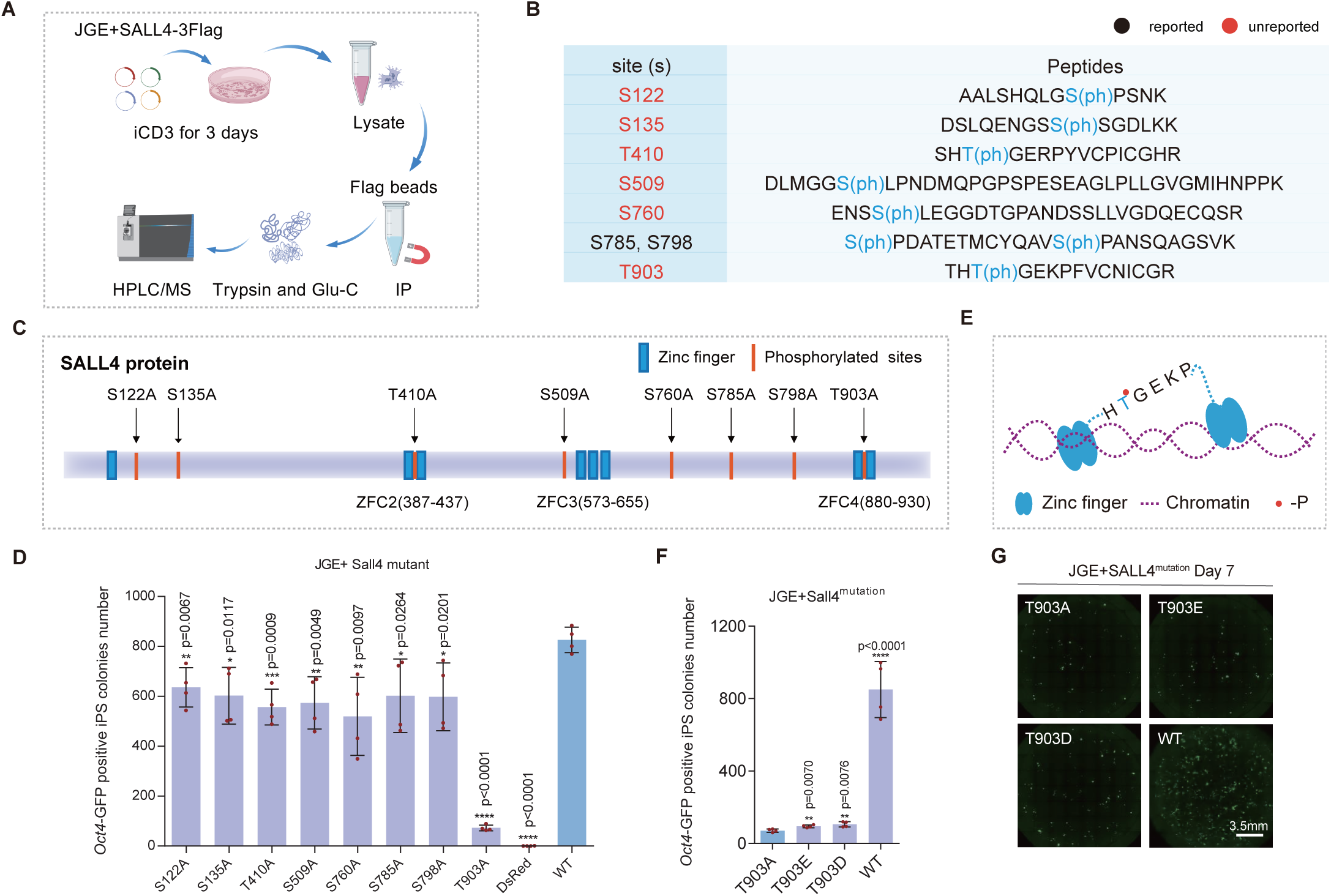
Phosphorylation at T903 is Critical for SALL4 Function. (A) The schematic diagram illustrates the data collection strategy for IP-MS in this figure. Created with BioGDP.com^80^. (B) The table displays the identified phosphorylation modifications and their corresponding peptide sequences. Black text indicates previously reported phosphorylation sites, red text denotes unreported modification sites. (C) The schematic diagram illustrates the relative positions of the identified sites within the SALL4 amino acid sequence in relation to the SALL4 zinc finger domains. (D) Bar plot for *Oct4*-GFP positive iPS colonies number of JGES reprogramming under each point mutation of SALL4 at Day 7, data are mean ± s.d., two-sided, unpaired t-test; n = 4 independent experiments, ****p < 0.001. (E) The schematic diagram shows the relative position of the HTGEKP motif within the ZFC4 linker of SALL4. (F) Bar plot for *Oct4-GFP* positive iPS colonies number of JGES reprogramming under each point mutation of SALL4 at Day 7, data are mean ± s.d., two-sided, unpaired t-test; n = 4 independent experiments, ****p < 0.001. (G) Images display the original well scanning results from panel (**e**), scale bar 3.5 mm.

We observed that T903 is located within the C2H2-type zinc finger domain ZFC4 (zinc finger complex 4), specifically on the linker sequence HTGEKP of SALL4^20^(Figure 1E). Given that aspartate (D) and glutamate (E) are commonly used as phosphomimetics for serine, threonine, and tyrosine due to their negatively charged carboxylate groups (-COO⁻), which mimic the phosphoryl group (-PO₄²⁻)^21^, we generated T903D and T903E mutants to assess functional mimicry. Surprisingly, neither mutant fully restored wild-type activity (Figure 1E and 1G), although both exhibited slightly higher reprogramming efficiency than the T903A mutant. These findings indicate that mere negative charge at T903 is insufficient to fully support SALL4 function, and suggest that the specific structural or electrostatic properties conferred by phosphorylation are necessary.

### T903A impairs chromatin opening and gene activation

The fact that T903 phosphorylation accounts for more than 90% of SALL4 activity encouraged us to focus on T903A for further analysis. We first collected bulk RNA-seq and ATAC-seq data at different time points (Figure 2A). PCA analysis based on the RNA-seq data shows that reprogramming induced by JGES^T903A^ proceeds more slowly than that induced by the JGES^WT^, with differences apparent as early as day 1 (Figure 2B). Further analysis reveal that the T903A seems to be deficient in gene activation compared to WT (Figure 2C). This suggests that pT903 may primarily be involved in gene activation. The heatmap and line chart of unsupervised clustering of genes between the JGES^WT^ and JGES^T903A^ groups, display the differential gene expression trends (Figure 2D and S2A), confirming the same observation in Figure 2C.

**Figure 2.**
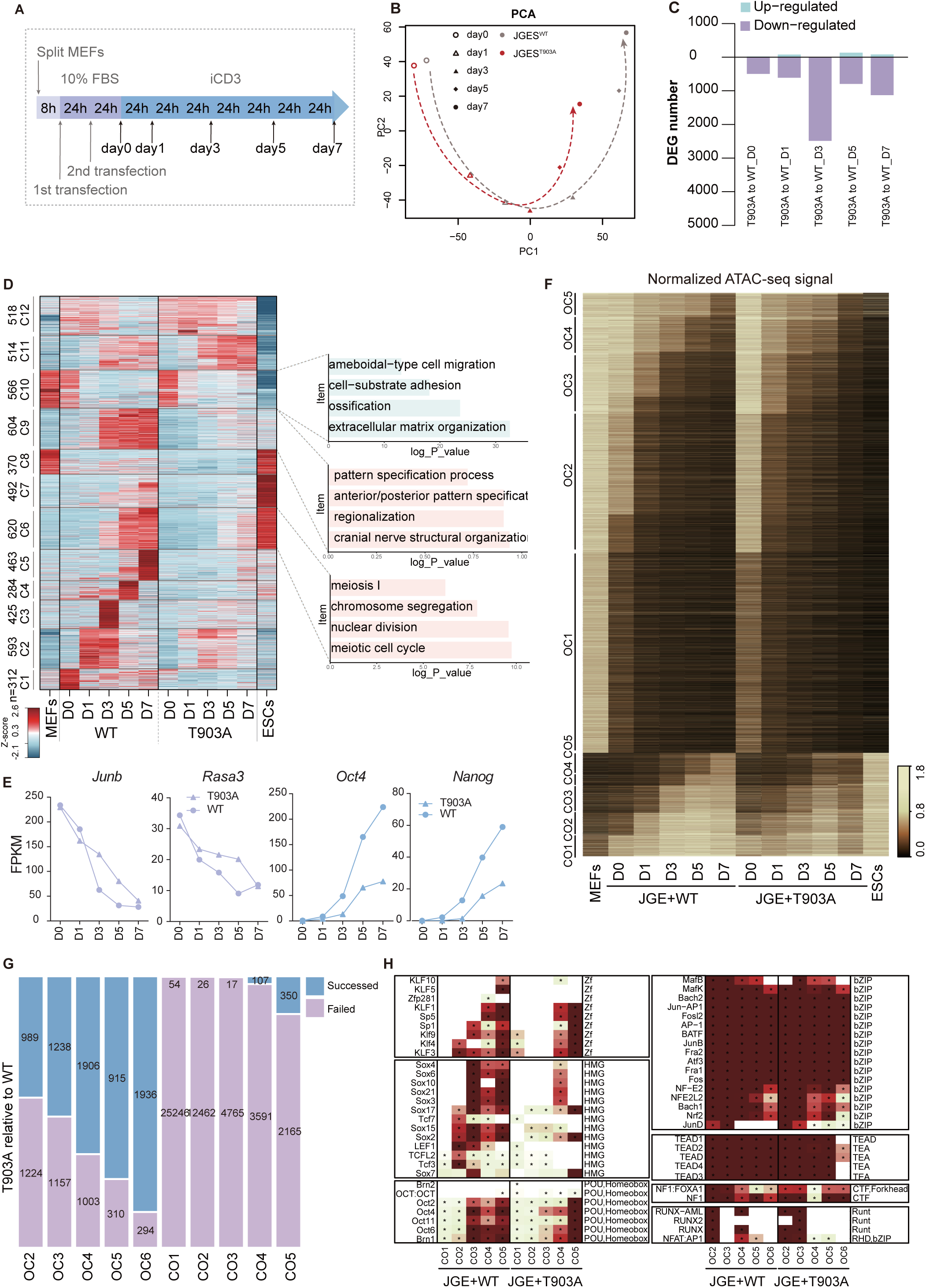
T903A Impairs Chromatin Opening and Gene Activation. **(A)** The flowchart illustrates the sample collection strategy for the RNA-seq and ATAC-seq experiments in this figure. (B) Transcriptome-based PCA illustrating trajectories of JGES^WT^ and JGES^T903A^ from Day 0 to Day 7. (C) Bar plot showing the number of up-regulated and down-regulated differentially expressed gene number in JGES^T903A^ compared to JGES^WT^. **(D)** Heat map showing the differentially expressed genes in JGES^WT^ and JGES^T903A^ reprogramming. Genes were clustered into 12 groups according to change pattern. The bar plot shows the gene ontology (GO) analysis of the groups. C1-12 indicate cluster 1-12. **(E)**The line chart illustrates the expression levels of individual genes in both JGES^WT^ and JGES^T903A^ groups. **(F)** Heat map showing the change in chromatin accessibility during reprogramming in JGES^WT^ or JGES^T903A^. Regions were categorized as either open-to-close (OC) or close-to-open (CO) events based on their temporal order in the control sample. **(G)** Bar plot showing the number of peaks with successful or failed chromatin state transitions in JGES^T903A^ compared to JGES^WT^ from Day 0 to Day 7. **(H)** Heat map showing motifs significantly enriched in each group. Motifs with at least 2-fold enrichment and a p-value < 0.01 are marked with asterisk.

Among these genes, C1–C7 and C9 represent gene sets that are upregulated in the JGES^WT^ group but show no or weaker upregulation in the JGES^T903A^ group, accounting for 63.24% of all differentially expressed genes. Specifically, cluster C6 consists of genes highly expressed in mESCs and significantly upregulated in the JGES^WT^ group, primarily associated with meiotic processes. In contrast, C9 comprises genes with relatively low expression in ESCs but significant upregulation in the JGES^WT^ group, mainly involved in the cranial nerve structural organization, regionalization and pattern specification process. In addition, the genes in C10, related to extracellular matrix organization, ossification, cell adhesion and migration, are effectively silenced in both groups. Among which, *Junb*^22^ and *Rasa3*^14^, two genes requiring suppression during the reprogramming. *Rasa3*, known to be directly regulated by the SALL4–NuRD axis, is silenced to comparable levels in both groups by day 7. In contrast, the pluripotency genes *Oct4* and *Nanog* are not sufficiently activated^23^ (Figure 2E).

Consistent with the RNA-seq results, ATAC-seq data further revealed that T903A specifically impairs gene activation, but not silencing function of SALL4 (Figure 2F, 2G, and 2H). During reprogramming, closed-to-open (CO) and open-to-closed (OC) transitions were defined as switch events and quantified as dynamic peaks^11^. T903A showed impaired opening but preserved closing relative to WT, suggesting activation-specific accessibility defects with minimal effect on chromatin closure.

### T903A impairs SALL4 and BAF interaction

Based on data above, T903A may potentially affect the interaction between SALL4 and gene-activating CFMs, like TET and BAF^13,24,25^. To test this idea further, we performed IP-MS both in JGES^WT^ and JGES^T903A^ at day1, as gene activation occurs as early as day 1 (Figure 2D). However, there are no TET related peptides detected in the IP-MS datasets at this stage, likely due to the low expression levels of TET RNA (S3A and S3B) and protein^26^. As expected, we show that SALL4-NuRD interaction are nearly identical between the WT and T903A ^27^ (Figure 3A, S3C and S3D), providing further evidence that T903A mutation has almost no effect on the gene silencing function of SALL4.

**Figure 3.**
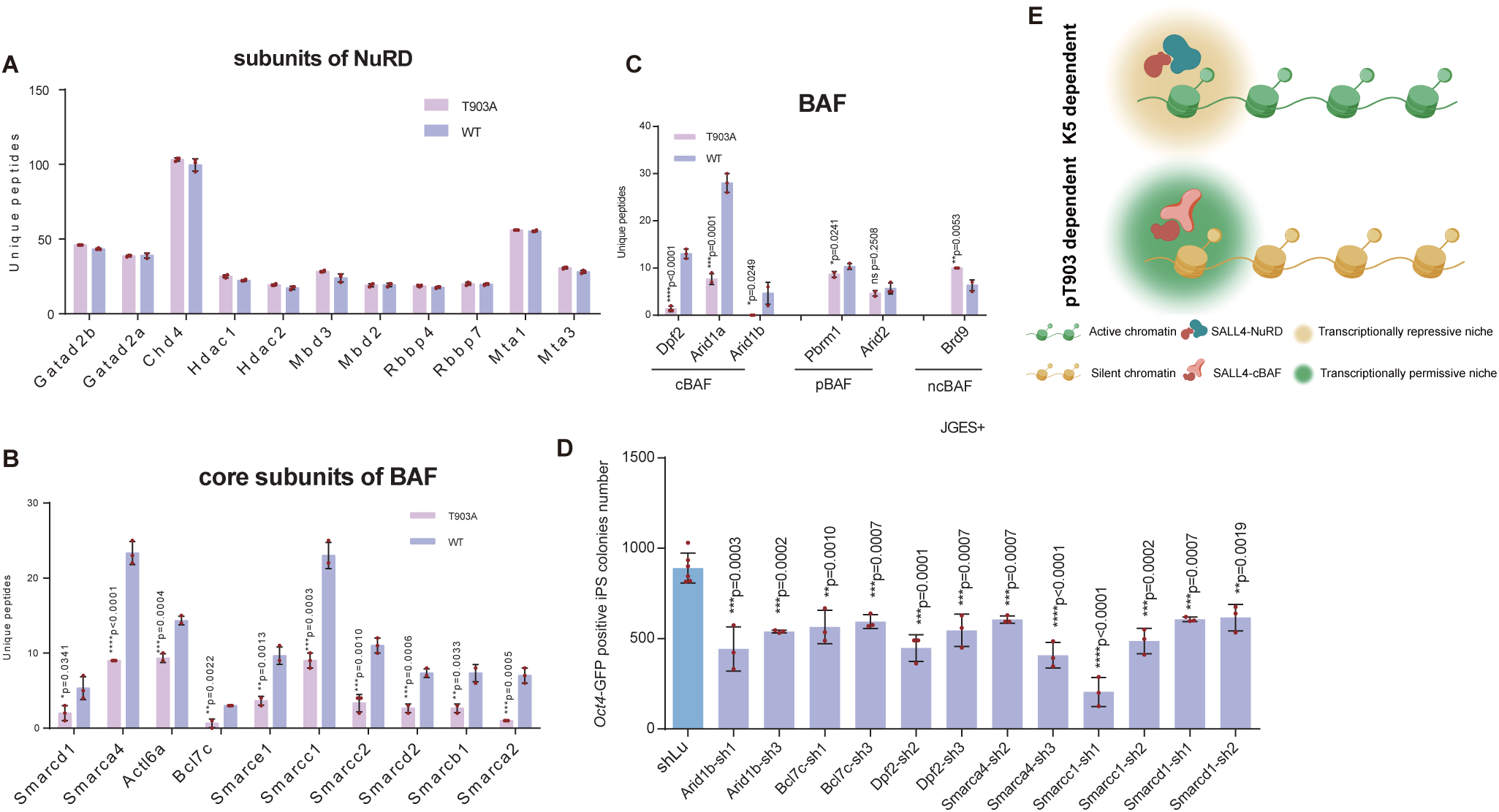
T903A Impairs SALL4 and BAF Interaction. (A) The bar chart shows the binding between SALL4 and each subunit of the NuRD complex at day 1 of JGES reprogramming both in the WT and T903A groups, data are mean ± s.d., two-sided, unpaired t-test; n = 3 independent experiments, ****p < 0.001. (B) The bar chart shows the binding between SALL4 and core subunits of the BAF complex at day 1 of JGES reprogramming both in the WT and T903A groups, data are mean ± s.d., two-sided, unpaired t-test; n = 3 independent experiments, ****p < 0.001. (C) The bar chart shows the binding between SALL4 and core subunits of cBAF, pBAF and ncBAF at day 1 of JGES reprogramming both in the WT and T903A groups, data are mean ± s.d., two-sided, unpaired t-test; n = 3 independent experiments, ****p < 0.001. (D) The bar chart shows the effect of knockdown of each BAF subunit on the number of *Oct4*-GFP positive iPSC colonies number in the JGES^WT^ reprogramming system. Data are mean ± s.d., two-sided, unpaired t-test; n = 3 independent experiments, ****p < 0.001. (E) The schematic diagram illustrates the interaction characteristics between SALL4 and NuRD or BAF complexes. Created with BioGDP.com^80^.

Surprisingly, there is a marked reduction of SALL4^T903A^-BAF interaction compared to that of SALL4^WT^(Figure 3B), suggesting that the loss of T903A gene activation may be associated with the reduced BAF interaction. We further show that cBAF (canonical BAF) is the only one markedly impaired compared to pBAF (polybromo-associated BAF) or ncBAF (non- canonical BAF) (Figure 3C). a control, we show that the RNA expression levels of BAF complex subunits remain largely similar between the WT and T903A groups (S3E and S3F), which excludes the impact of decreased BAF expression on the SALL4^T903A^-BAF interaction. Knockdown of each subunit of the BAF complex significantly inhibited the reprogramming efficiency of JGES^WT^ (Figure 3D and S3G), which is consistent with the decreased reprogramming efficiency caused by JGES^T903A^.

The BAF complex lacks DNA sequence specificity and relies on transcription factors for chromatin targeting^28^. It has been reported that SALL4 interacts with SMARCD1/BAF60a^25^ and DPF2^29^, two subunits of the BAF complex. To gain insight into the potential interactions between SALL4 and BAF, we performed molecular docking experiments^30,31^. The results show that SMARCD1 interacts with the phosphorylated T903 residue on SALL4 via its K156 and R163 residues (S3H), DPF2 interacts with the phosphorylated T903 residue on SALL4 via its V48, A49, Q50 and S51 residues (S3I). We also simulated the interactions between pT903 and both SMARCC1 and ARID1B, and the results showed no direct binding with either protein (data not shown). The interaction of SALL4-SMARCD1 and SALL4-DPF2 are abolished upon dephosphorylation or mutation of T903 to alanine, aspartic acid, or glutamic acid (S3H and S3I), providing further evidence supporting the results we observed for the failure of constitutively phosphomimetic residues (D/E) to substitute for pT903 in rescuing reprogramming (Figure 1E and 1F).

We propose that SALL4 recruits cBAF to repressive chromatin through pT903 site, establishing a transcriptionally permissive niche; the NuRD complex to active chromatin through K5 site, establishing a transcriptionally repressive niche (Figure 3E).

### BMP4 signals to SALL4 via kinases and phosphatases

The T903 phospho-switch for SALL4-BAF axis provides an ideal model to dissect their upstream signaling regulators. We have previously shown that BMP4 inhibits SALL4-mediated reprogramming^32^, thus, in light of our current findings, raising the possibility that BMP4 inhibits SALL4 by upregulating a phosphatase or downregulating a kinase targeting the pT903 site.

To search for such candidates, we intersected the genes up- or down-regulated genes under BMP4+/− conditions with known phosphatase and kinase gene sets (Figure 4A and Supplementary Table 1-2). Specifically, A1 represents phosphatases upregulated upon BMP4 treatment, A2 denotes kinases upregulated after BMP4 exposure, A3 indicates phosphatases downregulated by BMP4, and A4 refers to kinases downregulated following BMP4 treatment. Figure 4b-c and Extended Data Figure 4A and 4B display the top 30 genes within each of these four gene sets. The screening results suggest that BMP4 may influence the phosphorylation modification of pT903 by regulating the expression levels of *Dusp9* (Dual Specificity Phosphatase 9) ^33^and *Pdk1* (Pyruvate Dehydrogenase Kinase 1)^34^.

**Figure 4.**
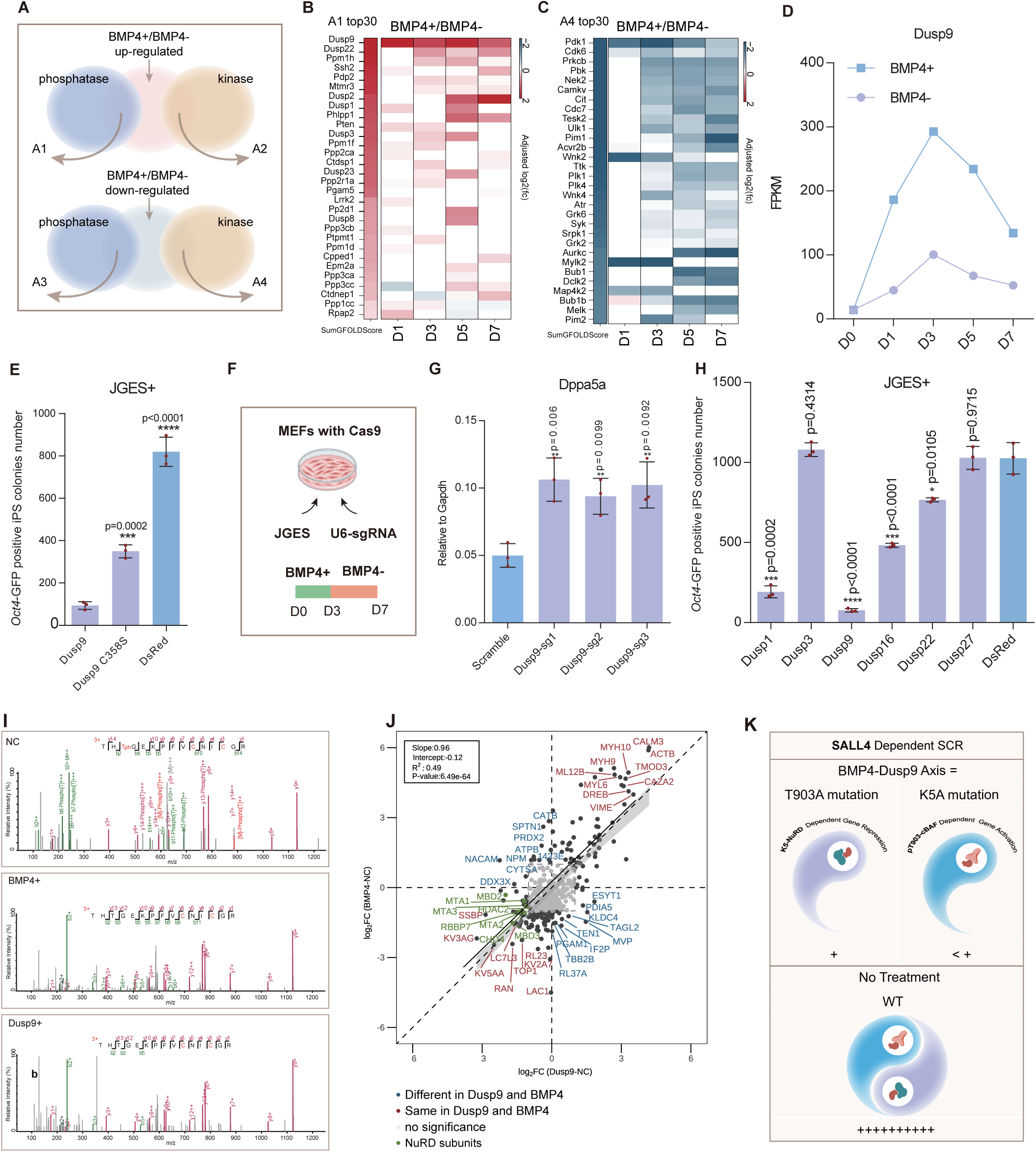
BMP4 Signals to SALL4 via Kinases and Phosphatases. (A) The Venn diagram illustrates the overlapping classification of genes between BMP4 treatment and the phosphatase/kinase gene sets. (B-C) The top 30 genes in A1 and A4 ranked by SumGFOLDScore, calculated as the sum of GFOLD values. (D) The line graph illustrates the activation of the Dusp9 by BMP4 treatment, based on time-course bulk RNA-seq data. (E) The bar chart shows the effect of over expression of different genes on the number of *Oct4*-GFP positive iPSC colonies number in the JGES reprogramming system. Data are mean ± s.d., two-sided, unpaired t-test; n = 3 independent experiments, ****p < 0.001. (F) The schematic diagram illustrates the knockdown strategy for DUSP9. (G) The histogram shows the relative expression levels of the late stage pluripotent gene *Dppa5a* in different groups. Data are mean ± s.d., two-sided, unpaired t-test; n = 3 independent experiments, ****p < 0.001. (H) The bar chart shows *Oct4*-GFP positive iPS colonies number of JGES reprogramming under over expression of different genes at Day 7, data are mean ± s.d., two-sided, unpaired t-test; n = 3 independent experiments, ****p < 0.001. (I) The image displays the IP-MS spectra of the peptide containing the T903 site under different treatments. (J) The quadrant plot illustrates the correlation of SALL4 interactome profiles between BMP4 treatment and DUSP9 over expression, fold change > 2, p-value < 0.05. (K) The schematic diagram illustrates the impact of the BMP4-DUSP9 axis on SALL4 during SALL4 dependent somatic cell reprogramming (SCR).

Our previous findings indicate that BMP4 exhibits nearly 100% inhibition in the JGES reprogramming system, with its inhibitory effect primarily occurring between day 1-3^32^. Notably, the activating effect of BMP4 on Dusp9 expression begins to decline after day 3, aligning precisely with its inhibitory window (Figure 4D). Meanwhile, overexpression of *Dusp9* inhibits JGES^WT^ reprogramming significantly, which dependents on the catalytic site, C358 ^33,35^ (Figure 4E). Moreover, knockdown of Dusp9 can rescue the inhibitory effect induced by BMP4 treatment during the first three days of reprogramming (Figure 4F, 4G, S4C, S4D and S4E). DUSP9 belongs to the MAP kinase phosphatase family and primarily functions to negatively regulate the activity of MAPK signaling pathways^36^. In addition to DUSP9, DUSP1, DUSP16, and DUSP22 also significantly inhibit reprogramming in the JGES system (Figure 4H), which suggests that the regulation of pT903 by the DUSP family may involve functional redundancy.

PDK1 is a key kinase in the PI3K/AKT signaling pathway through phosphorylating AKT at Thr308, thereby regulating cell survival, proliferation, growth, and apoptosis^34^.The transcriptional regulation of PDK1 by BMP4 is opposite to that of DUSP9, and knockdown of PDK1 significantly inhibits JGES reprogramming (S4F, S4G and S4H). To corroborate this result, we performed kinase prediction^19^ for the peptide THT(p)GEKPFVCNICGR, and show that PDK1 is ranked 25^th^ (Supplementary Table 3), suggesting that it is a plausible candidate for pT903.

To verify the association between BMP4-DUSP9 axis-mediated inhibition of JGES and pT903 dephosphorylation, we performed SALL4 IP-MS on samples collected on day 3 with BMP4 treatment group, the DUSP9-overexpressing group, and the control group. The MS spectra and statistical analysis showed that phosphorylation at the T903 site was detected exclusively in the control group (Figure 4I, S4I and S4J). These data provide direct evidence linking the BMP4-DUSP9 axis to the dephosphorylation of pT903. Using IP-MS interactome profiling (Figure 4J and Supplementary Table 4), we found that BMP4 treatment and DUSP9 overexpression produce highly consistent alterations in the SALL4 interaction proteome (slope: 0.96). Furthermore, overexpression of DUSP9 also disrupts the interaction between SALL4 and the NuRD complex (marked in green). The mass spectrometry results indicate that the BMP4-DUSP9 axis inhibits the reprogramming capacity of SALL4 through a dual mechanism: SALL4^pT903^ dephosphorylation and SALL4-NuRD dissociation (Figure 4K).

### T903A causes severe developmental defects in mice

Somatic cell reprogramming approximates the reverse of normal development. To investigate the critical role of pT903 during development, we first introduced the T903A point mutation into mouse embryonic stem cells using prime editing mediated by single-stranded oligodeoxynucleotides (SSODNs)^37^ (S5A and S5B), followed by tetraploid complementation to assess the impact of this point mutation on mouse embryonic development^38^ (Figure 5G). When cultured in serum + 2i/L medium for stemness maintenance^39^, mESCs with a homozygous T903A mutation displayed a distinct, flattened colony morphology with significantly increased intercellular spaces (Figure 5A). This contrasts with the dome-shaped colonies of wildtype mESCs and is reminiscent of human primed state embryonic stem cells^40^. Despite the distinct morphological alterations, both the proliferation rate (Figure 5B) and the transcript levels of pluripotency-associated genes (Figure 5C) in mutant ESCs remained comparable to those in wild-type controls, which is consistent with a recent study showing that morphological changes do not necessarily compromise the stemness of mouse embryonic stem cells^41^. In contrast to the strong suppression of large-scale gene activation during reprogramming caused by the point mutation, up regulated genes (n=638) slightly outnumbering down regulated genes (n=461) in mutant ESCs compared to the wild-type group (Figure 5D). Among them, T903A mutation resulting in significant up regulation of the two crucial imprinted genes, *Meg3* and *Igf2,* which are related to stem cell differentiation.^42–44^ Besides, the Gene Ontology (GO) Analysis indicate SALL4^T903^ participates in silencing of development-related genes, which is consistent with previously reported phenotype in SALL4 knockout mESCs (Figure 5E). Consistent with the GO enrichment analysis of C6 in Figure 2d, the T903A point mutation in ESCs also leads to the down regulation of genes associated with meiosis (Figure 5F), Notably, SALL4 has been reported to co-localize with ZBTB16 and SOX3 at over 12,000 promoters associated with the regulation of meiosis. Our data suggest that SALL4 mediated regulation of differentiation and mitosis-related genes could be regulated through phosphorylation at the T903 site.

**Figure 5.**
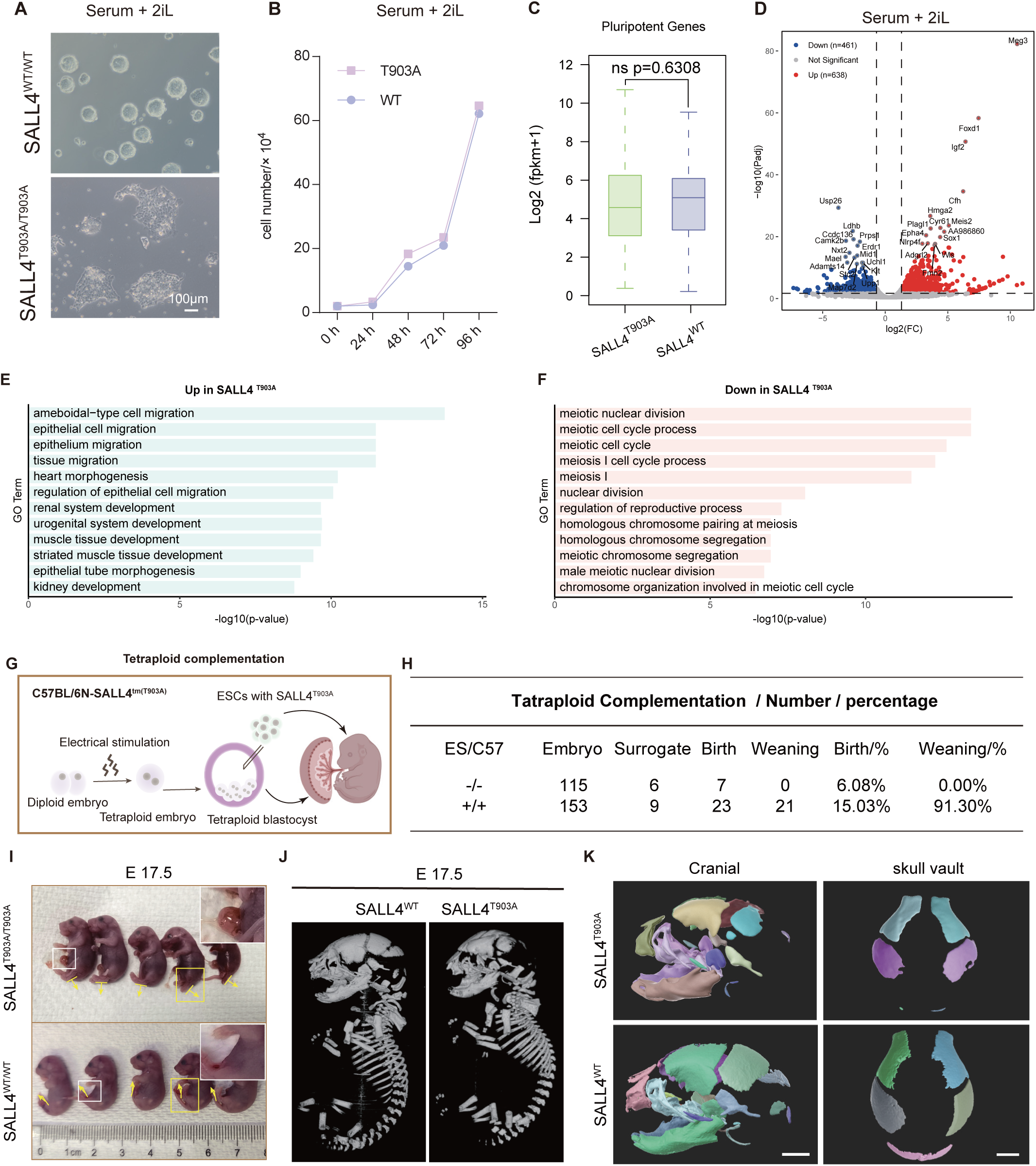
T903A Causes Severe Developmental Defects in Mice. **(A)**Bright-field images show the morphological features of SALL4^WT^ and SALL4^T903A^ homozygous mutant embryonic stem cells cultured in Serum + 2i/L condition. Scale bar = 100 μm. **(B)** The line graph shows the cell number of SALL4^WT^ and SALL4^T903A^ homozygous mutant embryonic stem cells across different time points, starting from an initial seeding density of 2×10⁴cells, data are mean ± s.d., two-sided, unpaired t-test; n = 3 independent experiments. **(C)** Box plot indicates the expression levels of pluripotency-associated genes between SALL4^WT^ and SALL4^T903A^ embryonic stem cells, two-sided, unpaired t-test; n=2 independent experiments. **(D)** Volcano plot shows the different expression genes between SALL4^WT^ and SALL4^T903A^ cells, fold change > 2, p-value < 0.05. Up-regulated transcripts are shown in red; down-regulated transcripts are in blue. **(E-F)** Functional enrichment analysis of genes up regulated and down regulated in SALL4^T903A^ compared to WT group. **(G)** The schematic diagram illustrates the strategy for tetraploid complementation assay. **(H)** The table presents the birth outcomes of mESCs from each experimental group. **(I)** Images show the external appearance of E17.5 mouse embryos from each experimental group. **(J)** Representative whole-body microCT reconstructions of the two mouse groups are presented, SkyScan 1276 Micro CT High resolution In-vivo Micro CT was used, 10 μm resolution. **(K)** 3D renderings of the skull and calvaria from each group of mice, processed using Amira software.

To further investigate the in vivo consequences, we injected wild-type (WT, +/+), and homozygous mutant (T903A/T903A, -/-) ESCs into tetraploid blastocysts. Cesarean sections were performed on surrogate dams at embryonic day 18.5 (E18.5) (Figure 5G). The WT group showed a birth rate of 15.03%, with a weaning rate of 91.30% among the newborns. In stark contrast, the birth rates for the mutant groups were markedly reduced to 6.08%, respectively. Furthermore, no pups from either mutant group survived past the weaning stage (Figure 5H), indicating that the T903A mutation leads to postnatal lethality. Previous studies have shown that SALL4 homozygous knockout mice die around E6.5-E7.5^45,46^. While mice homozygous for a truncation in the ZFC4 domain die around E10.5, heterozygous mice for these alleles are viable and can produce heterozygous offspring^47^. To explore the specific developmental defects in detail, we performed cesarean sections one day earlier, E17.5 (Figure 5I). The results showed no significant difference in body size or skin color between wild-type and mutant pups. However, the mutant pups exhibited a severe umbilical hernia (4/5 of T903A mutants vs. 1/5 of WT), a phenotype of abdominal wall hypoplasia.^48^ This was accompanied by a severe clubfoot phenotype (indicated by the golden arrow) (5/5 of T903A mutants vs. 0/5 of WT).^49^ These concurrent phenotypes—umbilical hernia and clubfoot—are consistent with a previously reported case of human developmental defects,^50^ this suggests that SALL4^T903^ site mutations could be incorporated into preconception genetic screening.^51^ Meanwhile, we fixed the bodies of wild-type and mutant pups with formalin and performed micro-computed tomography (microCT) scanning to visualize their skeletal structures (Figure 5J).^52^ The 3D reconstruction results revealed that the point mutation group exhibited significant cranial hypoplasia and a characteristically flattened skull vault (Figure 5K), which are likely to cause increased intracranial pressure, leading to potential brain herniation and central respiratory failure.^53–55^ This may constitute one of the primary causes for mutant mice to fail to survive after weaning. Furthermore, a severe spinal curvature (scoliosis, 1/5 of T903A mutants vs. 0/5 of WT) was noted in one of the mutant subjects (S5C). ^56^ These results suggest that *Sall4* plays critical roles in the development of multiple organs or tissues, especially those involving bone (Figure 5D).

## DISCUSSION

Based on the evidence presented above, we propose a phospho-switch for cell fate control. While phosphorylation and dephosphorylation have long been implicated in on/off control in signal transduction studies, a clear demonstration of their role in cell fate control has not been lacking. The only relevant exception is perhaps tumorigenesis, due to the fact that many proto-oncogenes are kinases and their constitutive activation leads to tumor formation^57^, as a result of cell fate mis-regulation. In most of the cases, the activated kinase accelerates cell cycle progression, which is distinct from cell fate change observed for SALL4 both in somatic cell reprogramming and mESCs stemness maintenance (Figure 2D and Figure 5F). Therefore, the phosphor-switch encoded by the HTGE motif in SALL4 is quite special in regulating precise cell fate control. Here are several considerations we can envision beyond our observations.

### The HTGE Motif

We performed a genome wide search and show that this switch motif also exists in 608 TFs in mice (Supplementary Table 5), include 1) ZBTB7a/b/c, which we have previously shown to be activated by BMP4 during mouse PNT (Primed-Naïve-Transition)^58^; 2) KLF4, whose function can be replaced by BMP4 in somatic cell reprogramming^59^; and 3) ZSCAN4C, a TF involved in totipotency that can be activated by BMP4 in mES cultured on feeder cells^60^, as examples. We notice with great interest that a consensus motif was initially reported in 1988 by Vogelstein and colleagues in GLI1^61^, a gene important for glioma and related cancer^62^. Like SALL4, GLI family of genes have been shown to be involved in cell fate determination during development^63^. As GLI TFs are at the final stage of hedgehog signaling, which mirroring our BMP-SALL4 axis, suggesting that this switch may function similarly but involving different sets of extracellular signals and intracellular partners. More work is needed to test these possibilities and define the mechanistic impact of this switch in both normal, disease and aging states ^64^. The apparent conservation of this motif in these HTGE TFs (TFs harboring the HTGE motif) implicated in cell fate control further support our overall hypothesis.

### The structure of HTGE Motif

Mouse SALL4 protein encodes the T903 motif as C2C12H3HT6C2C12H3H. T903 is sandwiched between two well conserved zinc figures, the type C2H2 (Figure 1E). In SALL4, it is called ZFC4 or zinc finger cluster 4, known previously to recognize AT-rich DNA. While ZFC4 has been implicated in cell fate control, T903 phosphorylation has not been reported and its role not identified. We speculate that the observed function of ZFC4 may be in part executed through T903 phosphorylation. Further studies are required to clarify this in the near future.

### Kinase and phosphatase for HTGE motif

We have tested several candidates for SALL4^pT903^ and presented evidence that DUSP9 is a likely candidate for pT903 dephosphorylation. We have also implicated PDK1 as the kinase for pT903. However, we cannot conclude here which ones are the canonical enzyme for pT903 modifications. Further studies are needed to clarify this. It is perhaps plausible to speculate that several kinases and phosphatases may be involved in SALL4 T903 phosphorylation and dephosphorylation. Clearly, for the other HTGE TFs, additional kinases/phosphatases may be tested under suitable conditions.

### HTG motif and BAF connection

We presented evidence that BAF is associated with SALL4 through pT903, thus, identifying an epigenetic downstream target for pT903. We have modeled this specific interaction and show that pT903 may interact through two BAF subunits, SMARCD1 and/or DPF2. We are continuing this line of investigation and hope to present physical evidence that this interaction is responsible for the observed cell fate control, thus, providing a mechanistic understanding of SALL4, especially pT903 in recruiting BAF complex and initiates transcription necessary for cell fate control as outlined in this report.

### Limitation of this study

We have not investigated other HTGE TFs. It is highly likely that these other TFs are subject to phosphorylation and dephosphorylation to control a specific fate during development or disease. Similarly, we have not characterized T903A knock-in model in depth enough. For instance, we have not identified the cell types affected by the T903A mutant embryos. We may need to perform single cell sequencing to identify the cell type affected which will further validate our hypothesis.

## RESOURCE AVAILABILITY

### Lead contact

All requests should be directed to the lead contact, Duanqing Pei.

### Materials availability

Materials will be available from the corresponding authors upon request.

### Data and code availability

- The ATAC-seq and RNA-seq data have been deposited in the Gene Expression Omnibus database under the accession number GSE316034.
- The mass spectrometry proteomics data have been deposited to the ProteomeXchange Consortium (https://proteomecentral.proteomexchange.org) via the iProX partner repository^65,66^ with the dataset identifier PXD072936.
- Code and any additional information required to reanalyze the data reported in this paper is available from the lead contact upon request.

## ACKNOWLEDGMENTS

We thank the faculty members of the Biomedical Research Core Facilities, high-Performance Computing Center and Laboratory Animal Resource Center of Westlake University. We also appreciate the help from all the members in Duanqing Pei’s lab. This work was supported by grants from The National Key Research and Development Program of China (2025YFA1804200), The National Natural Science Foundation of China (32500699), The Project funded by China Postdoctoral Science Foundation (2024M762944), The Zhejiang Provincial Natural Science Foundation of China (LZ25C060003), The Yangtze River Delta Sci-Tech Innovation Community Joint Research Project (2022CSJGG1000), The “Pioneer” and “Leading Goose” R&D Program of Zhejiang (2025C01115).

## AUTHOR CONTRIBUTIONS

J.M. designed and performed the main experiments; X.L., Z.J., and P.G. performed molecular cloning, qPCR and other verification experiments; C.Z., W.S. and Y.Y. analyzed the bioinformatic data; W.S. and Y.S. performed molecular docking; J.L, S.W. and H.Z. cultured mouse and prepared MEF cells; S.W. and Y.L. captured and reconstructed the microCT data; Y.C., S.L., R.S. and X.Z. performed and analyzed the IP-MS data; Y.F. provided the culture consultation of mES cells; B.W. provided the reprogramming system; D.P. supervised and conceived the whole study, wrote the manuscript, and approved the final version.

## DECLARATION OF INTERESTS

The authors declare no competing interests.

## STAR★METHODS

### ANIMALS

*Oct4*-DE-GFP (*OG2*-GFP) reporter-allele-carrying male mice (CBA/CaJ x C57BL/6J) were obtained from The Jackson Laboratory (004654). Rosa26-Cas9-EGFP homozygous male mice were obtained from the Laboratory Animal Center of Westlake University. The wild-type female 129 mice (129S2/SvPasCrl) were purchased from Vital River Laboratory Animal Technology Co., Ltd (Beijing). All animals were individually housed under the following conditions: a 12 h light/dark cycle, an ambient temperature of 20–26 °C, and 40%–70% humidity. Food and water were provided ad libitum. The animal studies were performed according to the applicable guidelines and regulations of the Institutional Animal Care and Use Committee of Westlake University (AP#25-130-PDQ-2), Hangzhou, China.

C57BL/6N-*Sall4*^tm(T903A)^ mice (-/-) were generated by Guangzhou Mingceler Biotech Co., Ltd. (Guangzhou, China). *Sall4*-targeting SSODNs (gcagccgcagccacgccgccaggccaagcagc actgctgcacacggtgtggaaagaacttctcgtctgccagtgccctgcagatccacgagcgaacacacgcgggagagaagcctttcgtgtgtaacatatgcgggcgggccttcaccacgaaaggcaacctgaaggtgggttccgacgggcgttggtgtgtgacgtctgt) with the mutation and sgRNA (ctcccgtgtgtgttcgctcg) were transfected into embryonic stem cells via electroporation to generate homozygous, (-/-), mutant cell lines. Knock in mice were subsequently generated with the Knock in cell lines through tetraploid embryo complementation. Animals were housed in plastic cages (2–3 mice per cage) under SPF (specific pathogen-free) conditions, with sentinel mice undergoing periodic pathogen screening throughout the study. Mice were housed under a 14-h light:10-h dark cycle, with temperature and relative humidity maintained at 23.5 ± 2.5°C and 52.5 ± 12.5%, respectively. They received standard laboratory chow and autoclaved water ad libitum. All experimental protocols were reviewed and approved by the Institutional Animal Care and Use Committee (IACUC) of Guangzhou Mingceler Biotech Co., Ltd.

### METHOD DETAILS

#### Cell lines

E13.5 mouse embryo fibroblasts (MEFs), regardless of sex, were generated by *OG2* mice and 129 mice, E13.5 mouse embryo fibroblasts with Cas9 (MEFs with Cas9), regardless of sex, were generated by Rosa26-Cas9-EGFP mice and 129 mice.After removing the integral organs, the tail, the limbs and head, the remaining tissues were cut into small pieces and digested (0.25% trypsin: 0.05% trypsin = 1:1; GIBCO) for 8-10 min at 37 °C. The isolated MEF cells were seeded on 0.2% gelatin (home-made, porcine skin) coated dish, cultured in fibroblast medium: DMEM-high glucose (Hyclone) contains 10% FBS (Vazyme, F103).

Plat-E cells were cultured in DMEM high-glucose media (Hyclone) supplemented with 10% FBS (Vazyme, F103).

Mouse embryonic stem cells (mESCs) were culture in DMEM high-glucose media (Hyclone) supplemented with 10% FBS (Gibco, a5669701), 1% GlutaMAX (GIBCO), 1% sodium pyruvate (GIBCO), 1% NEAA (GIBCO), 0.1 mM 2-mercaptoethanol (GIBCO), 1000 U/ml LIF (Millipore, ESGE107), 3 mM CHIR99021 (Sigma, SML1046), and 1 mM PD0325901 (Sigma, PZ0162).

All the cell lines have been confirmed as mycoplasma contamination-free with the Kit from Lonza (LT07-318).

#### DNA constructs

pMXs plasmids, pSuper-shRNA plasmids, pSuper-sgRNA plasmids were used for in vitro over-expression. The pMXs (retrovirus vector) were regularly used. The pSuper-shRNA plasmid (retrovirus vector) is commonly used for gene knockdown. Whereas, the pSuper-sgRNA plasmid was constructed by replacing the original shRNA in the pSuper-shRNA backbone with an sgRNA-scaffold cassette downstream of the U6 promoter. The sequences of sgRNA and shRNA used in this article are shown in Supplementary Table 6.

#### iPSCs generation

Plat-E cells were seeded in 10-cm dishes (7.5–8.5 × 10⁶ cells/dish) 12–16 h before transfection. For each dish, 10 μg plasmid DNA was mixed with 40 μL PEI (1 mg/mL) in 1 mL Opti-MEM (GIBCO, 31985070), incubated for 10–15 min, RT, and added to cells in 9 mL fresh medium (10% FBS/DMEM). The medium was replaced at 10–16 h post-transfection. Retrovirus (Plat-E) supernatant was collected at 48 h and 72 h. All supernatants were filtered (0.45μm) and stored at room temperature for ≤48 h.

*OG2* MEFs were seeded into a 24-well plate (1.5 × 10⁴ cells/well). Retrovirus (*Jdp2*: *Glis1*: *Esrrb*: *Sall4* = 2:1:1:2) was mixed with polybrene (4μg/mL) and Y27632 (5μM) for infection. Two rounds of retroviral infection were performed at a 24 h interval. Two days post-infection, medium was changed to iCD3 (DMEM-high glucose contains TV, VC, CHIR-99021, bFGF, mLIF, SGC0946, GSK-LSD1-2HCL, Y27632. 1% sodium pyruvate (GIBCO), 1% non-essential amino acids (GIBCO), 1% GlutaMAX (GIBCO), 0.1mM 2-mercaptoethanol (GIBCO), N2 (GIBCO), B27 (GIBCO)), refreshed daily. GFP⁺ colonies were imaged (Keyence, BZ-X810) and counted using ImageJ Particle Analysis.

#### Immunopurification-mass spectrometry

Cells were trypsinized, washed with cold PBS, and pelleted. Pellets were flash-frozen in liquid nitrogen and stored at –80°C. Whole-cell extracts were prepared by lysing pellets in ice-cold lysis buffer (50 mM Tris pH 7.4, 150 mM NaCl, 10% glycerol, 1% NP-40, 1 mM EDTA) supplemented with 1× Complete Protease Inhibitor Cocktail (Sigma, 1187358001), 1% PMSF (in isopropanol), and 100× phosphatase inhibitor (Yuanye Bio-Technology, R32811). Lysates were incubated on ice for 20 min, then rotated at 4°C for 1 h. Cleared lysates were obtained by centrifugation at maximum speed for 15 min at 4°C. The supernatant was incubated with anti-FLAG magnetic beads (Thermo Fisher, A36797) overnight at 4°C with rotation. Beads were washed 10 times with wash buffer (50 mM Tris pH 7.4, 150 mM NaCl, 10% glycerol, 0.01% NP-40, 1 mM EDTA) on ice. After removing the final wash, bound proteins were eluted by boiling in loading buffer (4% SDS, 10% 2-mercaptoethanol, 20% glycerol, 0.004% bromophenol blue, 0.125 M Tris pH 6.8) at 99°C for 10 min. Samples were stored at –80°C and avoid freeze-thaw cycle.

The IP samples were separated by SDS-PAGE gel, followed by in-gel digestion. Gel pieces were destained with 50 mM NH₄HCO₃ in 50% acetonitrile, dehydrated with 100% acetonitrile, then reduced with 10 mM DTT at 56°C for 60 min. After dehydration, alkylation was performed with 55 mM iodoacetamide in the dark for 45 min. Gel pieces were washed with 50 mM NH₄HCO₃, dehydrated, and rehydrated on ice with 10 ng/μl trypsin in 50 mM NH₄HCO₃ for 1 h. Digestion proceeded overnight at 37°C. Peptides were extracted with 50% acetonitrile/5% formic acid, followed by 100% acetonitrile, dried, and resuspended in 2% acetonitrile/0.1% formic acid. Peptides were loaded onto a reversed-phase column (15 cm × 75 μm) and separated using a gradient from 6% to 23% solvent B (0.1% formic acid in 98% acetonitrile) over 16 min, 23% to 35% in 8 min, increased to 80% in 3 min, and held at 80% for 3 min at 400 nl/min. MS was performed on a Q Exactive Plus coupled to an EASY-nLC 1000. Full scans (m/z 350–1800) were acquired at 70,000 resolution, followed by up to 20 MS/MS scans (NCE 28) at 17,500 resolution. Dynamic exclusion was set to 15 s, and AGC was 5×10⁴.

The MS/MS data were searched using Proteome Discoverer 2.4 with the following parameters: the target protein sequence database was used; enzyme specificity was set to Trypsin and Glu-C; up to two missed cleavages were allowed for phosphorylated peptides; the mass tolerance for precursor ions was 10 ppm, and for fragment ions was 0.02 Da; fixed modification was set as carbamidomethylation of cysteine; variable modifications included oxidation of methionine, acetylation of protein N-termini, and phosphorylation of serine/threonine/tyrosine; peptide ion score threshold was set above 20, and identification results were filtered at High peptide confidence.

#### RNA-seq data analysis

RNA-seq reads were processed using Trim Galore (v0.6.4) in paired-end mode to remove adapter sequences^67,68^. Trimmed reads were then aligned to the mm10 reference genome using HISAT2 (v2.2.1) in paired-end mode^69^. Gene and repeat locus expression levels were quantified using GFOLD (v1.1.4) with the gfold count module^70^. Differentially expressed genes in Figure 2c and Figure 4b, c were identified using the gfold diff module. Genes with |GFOLD value| > 1.5 were considered significantly differentially expressed. For Figure 5c, differential expression analysis was performed using DESeq2 (v1.38.3) based on raw read counts,^71^ with significance defined as adjusted p-value < 0.05 and |log₂(fold change)| > 1. Principal component analysis was conducted using the prcomp function in R (v4.2.3) on FPKM values. Unsupervised clustering of genes was performed using the k-means algorithm implemented in the amap package (v0.8.20) in R, with FPKM values as input, employing Euclidean distance, iter.max = 100, and nstart = 50. For visualization, the expression profiles of the clustered genes were z-score normalized across samples to highlight relative expression patterns. The list of phosphatases and protein kinases was obtained from the Gene Ontology database (http://amigo.geneontology.org/amigo). The list of pluripotency-associated genes was compiled from the paper by Shen et al.^72^

#### ATAC-seq data analysis

ATAC-seq reads were trimmed using Trim Galore (v0.6.4) and then aligned to the mm10 reference genome using bowtie2 (v2.4.5) in paired-end mode.^73^ The repetitive, low sequencing quality (mapq < 30) and mitochondrial DNA aligned reads were excluded using SAMtools (v1.16.1).^74^ In order to make the data comparable between different sequencing depths, the signals were normalized to one million reads for each sample, and the value were further compressed into a binary format (bigWig) for downstream analysis and data visualization. Peak calling was performed using MACS (v1.4.2) with parameters as follows: -g mm --keep-dup all --nomodel --shiftsize 25.^75^

Regions with Dynamic change in chromatin accessibility were identified by temporal changes in peak occupancy across time-series ATAC-seq samples. Regions gaining chromatin accessibility (close-to-open, CO) or losing accessibility (open-to-close, OC) were classified based on the temporal pattern of peaks being detectable. Normalized ATAC-seq signal were extracted for each region using bigWigSummary (UCSC tools) and visualized as heatmaps.^76^

#### Motif analysis

Motif enrichment analysis was performed using HOMER (v4.11.1) on differentially accessible regions identified from ATAC-seq peak data.^77^ The findMotifsGenome.pl command was executed with the parameters -mask to mask repetitive elements during the search. HOMER used hypergeometric test to determine the motif enrichment and also test the similarity between the motif we identified to known factors. Only motifs with a -log₁₀(p-value) > 2 and an enrichment score > 2 are displayed in the plot.

#### Gene Ontology analysis

Functional annotation was performed using the clusterProfiler (v4.6.2).^78^ Gene Ontology terms for each functional cluster were summarized to a representative term, and adjusted p-values were plotted to show the significance.

#### Molecular docking

Interactions between SALL4 (including its variants) and SMARCD1 were predicted using AlphaFold3 web server (EMBL-EBI, https://alphafoldserver.com).^31^ The amino acid sequences of both proteins were retrieved from UniProt^30^. The resulting complex structures were visualized and analyzed using PyMOL (v3.1.6.1) to characterize interaction interfaces.^79^

#### Statistics

All statistical analyses were performed using GraphPad Prism or R. The details of the statistical tests applied are provided in the figure legends.

## Notes

### Competing Interest Statement

The authors have declared no competing interest.

