## Supplemental Figure 1 for "A Phospho-Switch for Cell Fate Control"

### Figure S1

**A**

**S122**

Sequence: AALSHQLGSPSNK, S9-Phospho (79.96633 Da)

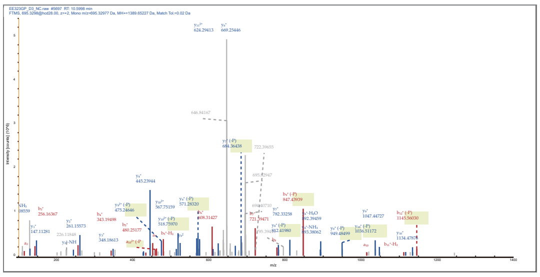

**S760**

Sequence: ENSSLEGGDTGPANDSSLLVGDQECQSR, C25-Carbamidomethyl (57.02146 Da) S4-Phospho (79.96633 Da)

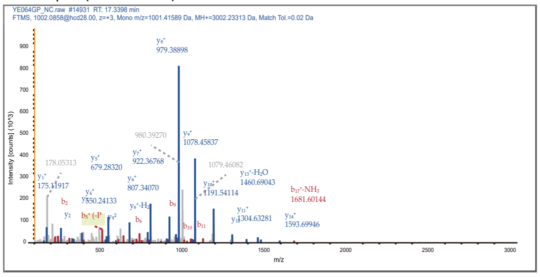

**S135**

Sequence: DSLQENGSSSGDLKK, S9-Phospho (79.96633 Da)

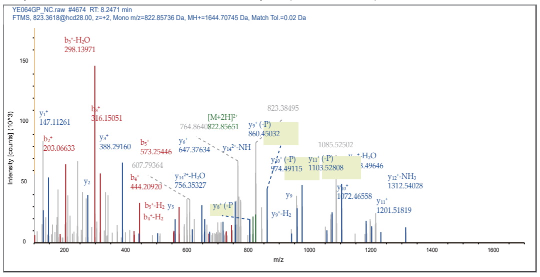

**S785**

Sequence: AALSHQLGSPSNK, S9-Phospho (79.96633 Da)

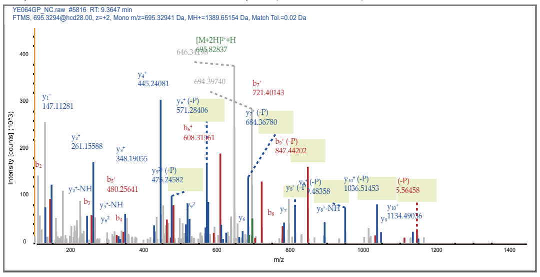

**T410**

Sequence: SHTGERPYVCPICGHR, C10-Carbamidomethyl (57.02146 Da), C13-Carbamidomethyl (57.02146 Da), T3-Phospho(79.96633 Da)

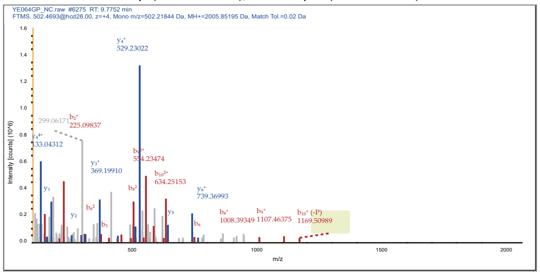

**S798**

Sequence: SPDATETMCYQAVSPANSQAGSVK, C9-Carbamidomethyl (57.02146 Da), S1-Phospho (79.96633 Da), S14-Phospho (79.96633 Da)

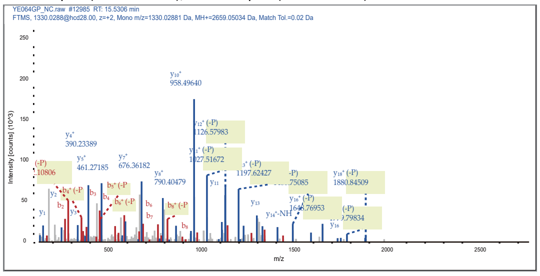

**S509**

Sequence: DLMGGSLPNDMQPGPSPESEAGLPLLGVGMIHNPPK, S6-Phospho (79.96633 Da), M3-Oxida (15.99492 Da), M11-Oxida (15.99492 Da), M30-Oxida (15.99492 Da)

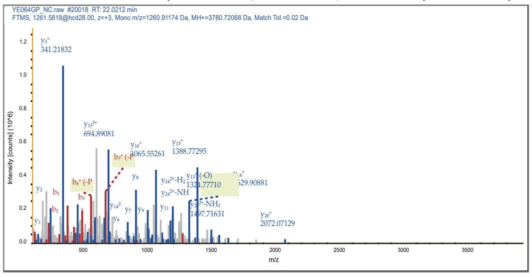

**T903**

Sequence: THTGEKPFVCNICGR, C10-Carbamidomethyl (57.02146 Da), C13-Carbamidomethyl (57.02146 Da), T3-Phospho (79.96633 Da)

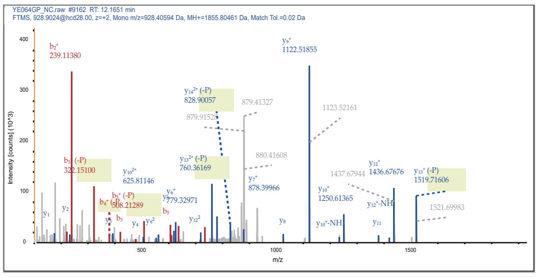

**Figure S1. Phosphorylation at T903 is critical for SALL4 function, related to Figure 1**

(A) Images display the secondary Mass spectra of the peptides containing each identified phosphorylation site shown in Figure 1B.
