## Supplemental Figure 2 for "A Phospho-Switch for Cell Fate Control"

**Figure S2**

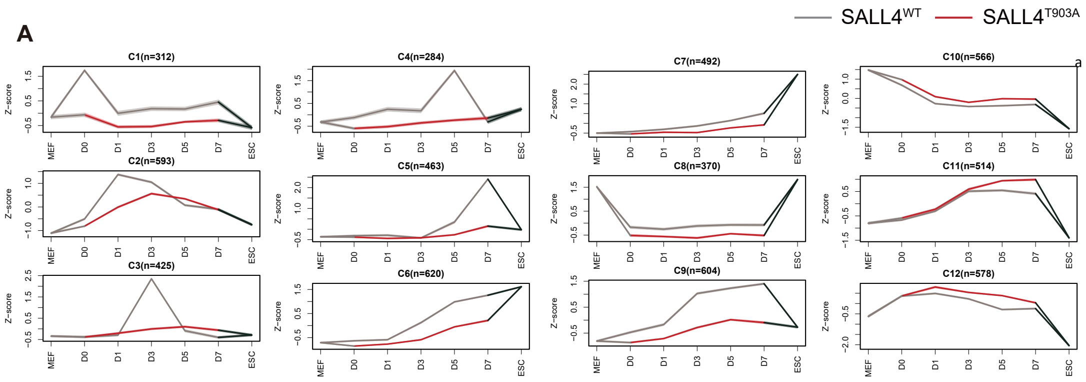


**Figure S2. T903A impairs chromatin opening and gene activation, related to Figure 2.**

(A) Line plots showing the expression dynamics of gene groups during reprogramming in $JGES^{WT}$ and $JGES^{T903A}$.
