## Supplemental Figure 3 for "A Phospho-Switch for Cell Fate Control"

### Figure S3

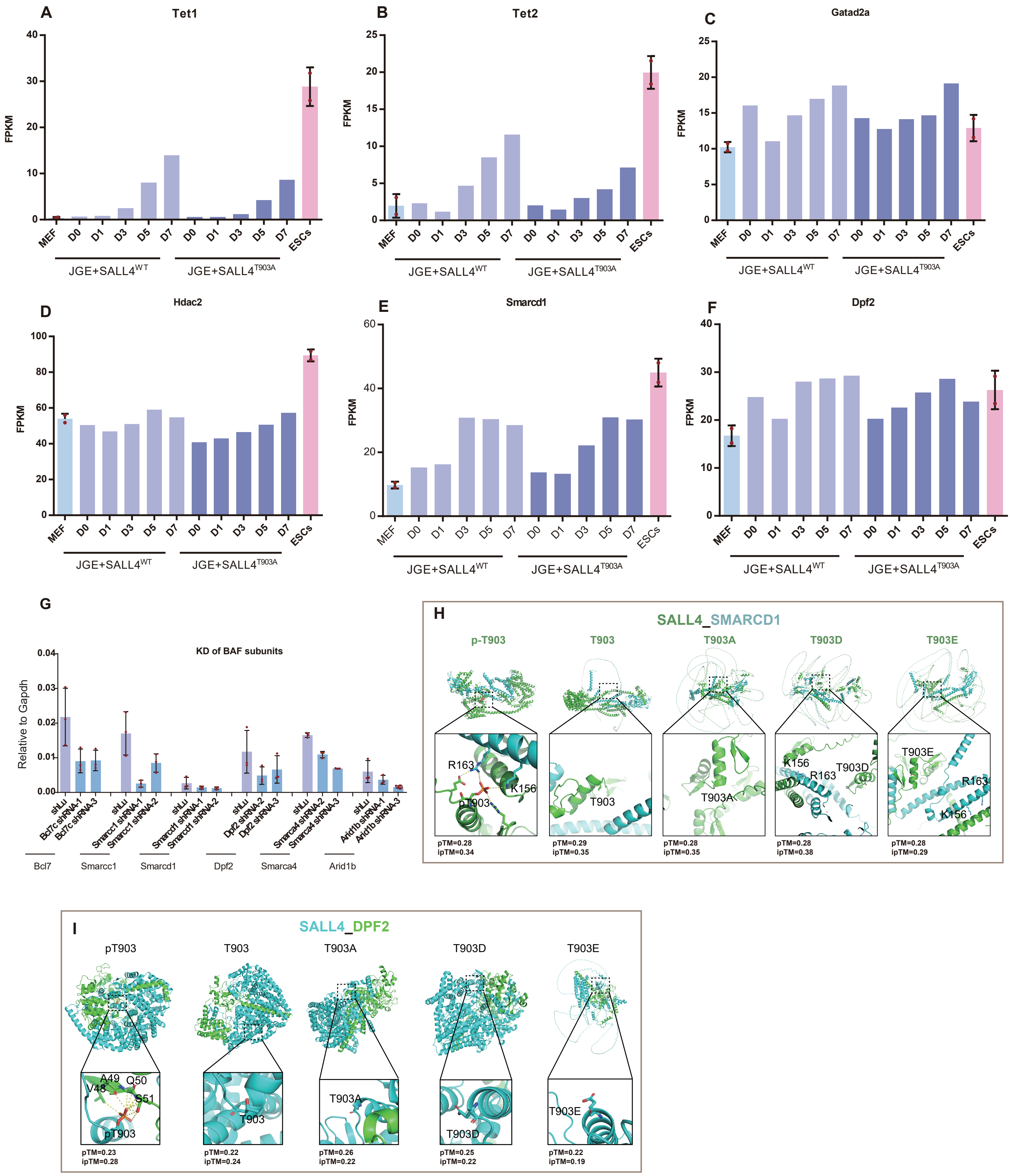


**Figure S3 T903A impairs SALL4 and BAF interaction, related to Figure 3.**
(A-F) The histogram displays the expression levels of genes in the indicated system from the Bulk RNA-seq results.
(G) The histograms shows the knockdown efficiency of the shRNAs targeting BAF subunits used in Figure 3D, data are mean ± s.d., two-sided, unpaired t test; n = 3 independent experiments.
(H-I) AlphaFold-Multimer predictions of SALL4–SMARCD1 and SALL4-DPF2 interaction modes and visualized and analyzed using PyMOL (v3.1.6.1). pTM (the predicted template model) and ipTM (the interface predicted template model) scores reflect the confidence in global fold and interface accuracy
