## Supplemental Figure 4 for "A Phospho-Switch for Cell Fate Control"

### Figure S4

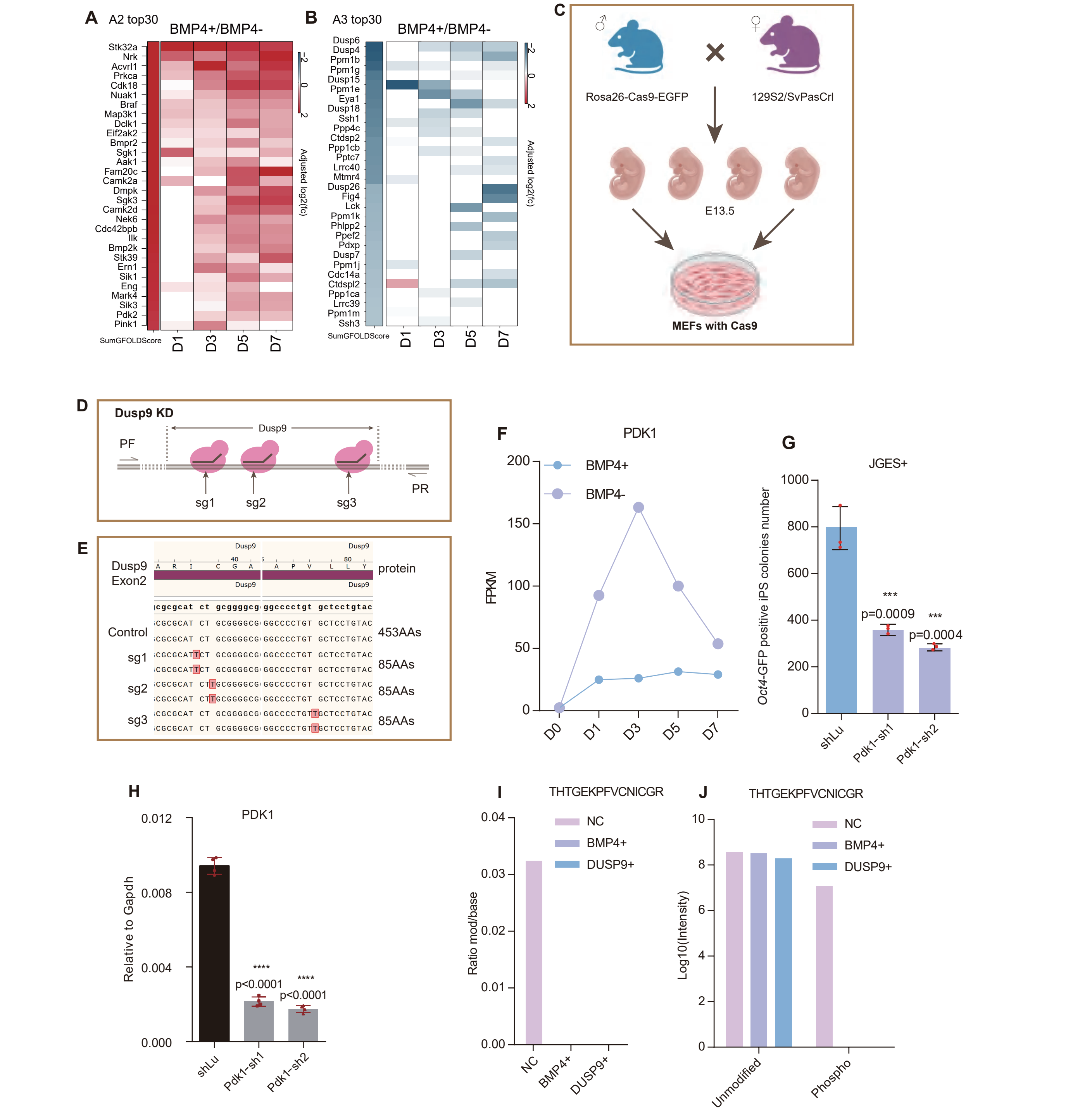


**Figure S4. BMP4 signals to SALL4 via kinases and phosphatases, related to Figure 4**
(A-B) The top 30 genes in A2 and A3 ranked by SumGFOLDScore, calculated as the sum of GFOLD values.
(C-D) The schematic diagram illustrates the preparation strategy for embryonic fibroblast cells (E13.5) derived from Cas9 transgenic mice.
(E) The image shows the point mutations induced in the Dusp9 genome by different sgRNA treatments. The results indicate that all three sgRNAs cause premature termination of DUSP9 translation, shortening the protein from 453 amino acids (WT) to 85 amino acids in the experimental groups, as visualized in SnapGene.
(F) The line graph illustrates the suppression of the Pdk1 by BMP4 treatment, based on time-course bulk RNA-seq data.
(G) The bar chart shows the effect of Pdk1 KD on the number of GFP-positive iPSC colonies in the JGES reprogramming system. Data are mean ± s.d., two-sided, unpaired t test; n = 2 independent experiments, ****p < 0.001.
(H)The histogram shows the knockdown efficiency of the shRNAs targeting Pdk1 used in panel.
(G) data are mean ± s.d., two-sided, unpaired t test; n = 3 independent experiments, ****p < 0.001.
(I-J) The histograms illustrate the effects of different treatments on the phosphorylation modification of the peptide THTGEKPFVCNICGR
