## Supplemental Figure 5 for "A Phospho-Switch for Cell Fate Control"

### Figure S5

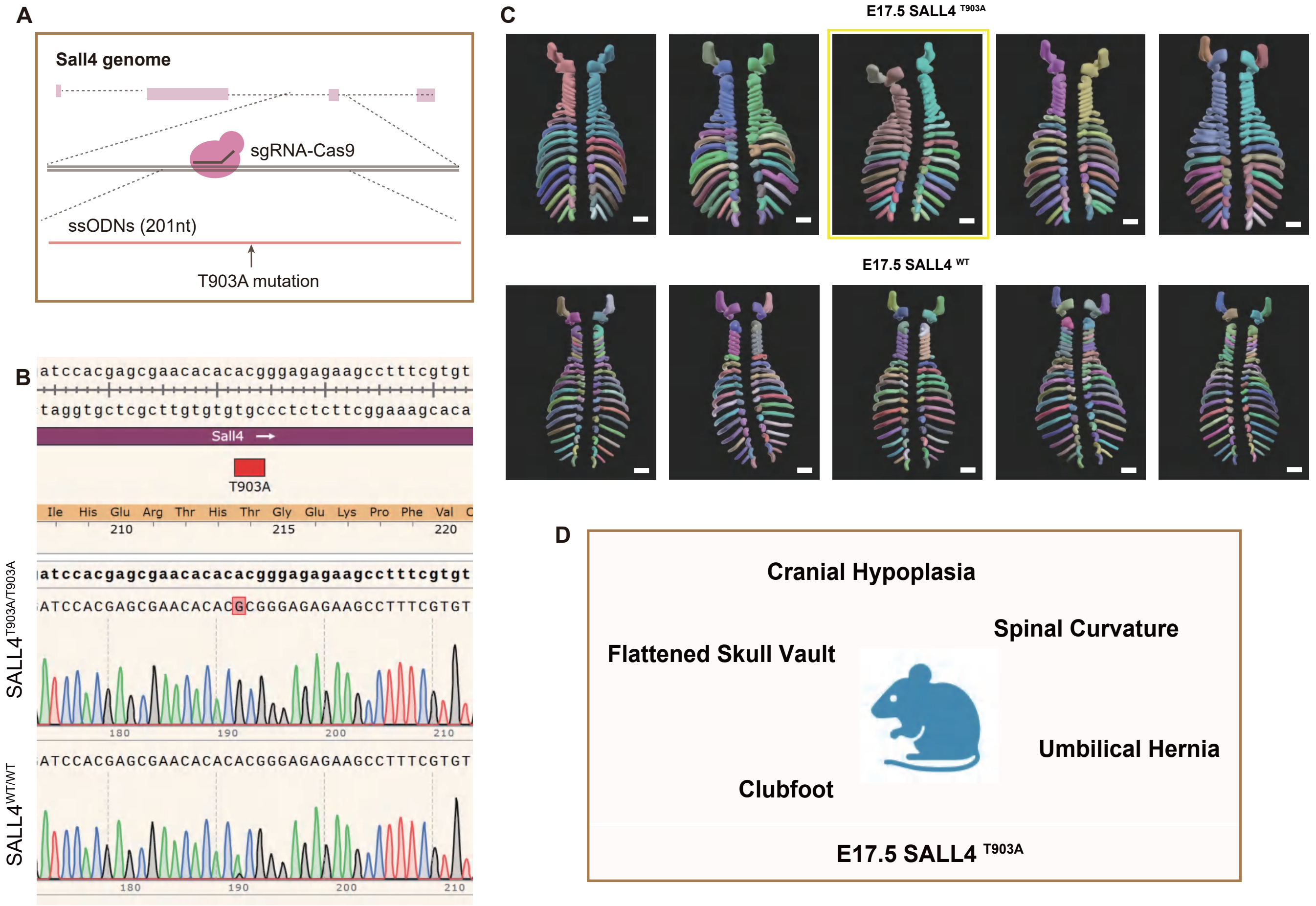


**Figure S5. T903A causes severe developmental defects in mice, related to Figure 5**
(A) Schematic of the gene-editing strategy for introducingthe SALL4 T903A point mutation.
(B) The genotyping results confirming the WT and homozygous mutant ESC lines are presented, as visualized in SnapGene.
(C) 3D renderings of the cervical vertebrae, thoracic vertebrae, and ribs from each group of mice, processed using Amira software. The golden box shows the severe scoliosis.
(D)The schematic illustrates the postnatal developmental defects observed in SALL4$^{-/-}$ mice.
